# Brief targeted memory reactivation during the awake state enhances memory stability and benefits the weakest memories

**DOI:** 10.1101/196436

**Authors:** Arielle Tambini, Alice Berners-Lee, Lila Davachi

## Abstract

Reactivation of representations corresponding to recent experience is thought to be a critical mechanism supporting long-term memory stabilization. Targeted memory reactivation, or the re-exposure of recently learned cues, seeks to induce reactivation and has been shown to benefit later memory when it takes place during sleep. However, despite recent evidence for endogenous reactivation during post-encoding awake periods, less work has addressed whether awake targeted memory reactivation modulates memory. Here, we found that brief (50ms) visual stimulus re-exposure during a repetitive foil task enhanced the stability of cued versus uncued associations in memory. The extent of external or task-oriented attention prior to re-exposure was inversely related to cueing benefits, suggesting that an internally-orientated state may be most permissible to reactivation. Critically, cueing-related memory benefits were greatest in participants without explicit recognition of cued items and remained reliable when only considering associations not recognized as cued, suggesting that explicit cue-triggered retrieval processes did not drive cueing benefits. Cueing benefits were strongest for items and participants with the poorest initial learning. These findings expand our knowledge of the conditions under which targeted memory reactivation can benefit memory, and in doing so, support the notion that reactivation during awake time periods improves memory stabilization.

## INTRODUCTION

After the initial encoding of events into memory, post-encoding processes are thought stabilize new experiences into long-term memory representations^1^. The reactivation of representations corresponding to recent experience, or ‘replay’, is a major candidate mechanism thought to support the memory stabilization^2–4^. In particular, the sequential reactivation of hippocampal ensembles during sharp-wave ripple events (SWRs), in conjunction with hippocampal-cortical interactions, are thought to be key mechanisms underlying the stabilization of representations of recent experience^5,6^. Reactivation and SWRs occur primarily during ‘off-line’ brain states, such as sleep and quiescent awake periods^7–9^.

Evidence for reactivation as a mechanism underlying memory stabilization comes from work showing that the strength of endogenous SWR-related activity or reactivation of encoding patterns is positively related to later memory^10–16^. The most compelling evidence for the functional relevance of off-line reactivation comes from work that causally manipulates reactivation events and examines their impact on memory^17,18^. One powerful approach, termed targeted memory reactivation, seeks to exogenously induce reactivation via sensory information or the re-exposure of a subset of cues from learned associations^19,20^. Cue re-exposure during non-rapid-eye-movement (NREM) sleep enhances memory for cued versus uncued associations^19,21–25^. This phenomenon, combined with evidence that cueing can influence hippocampal activity and the content of reactivated representations^26,27^, provides strong evidence linking reactivation with memory.

The majority of studies to date using targeted memory reactivation have performed cueing during NREM or other phases of sleep^23,28^. Although it is widely acknowledged that sleep is an important time period for memory consolidation and reactivation^29^, awake periods may also contribute to memory consolidation. It has been proposed that reactivation and SWRs occur during time periods when the hippocampus is not engaged in encoding novel information^7,30,31^. In rodents, hippocampal reactivation and SWRs clearly occur during the awake state, particularly during quiescence or pauses in ongoing behavior^32–34^. Evidence for memory-related hippocampal reactivation, connectivity, and SWR activity in humans has been found during post-encoding awake periods^10,13–16,35,36^. Moreover, behavioral work has shown that the presence of rest periods after encoding can benefit later recall or associative memory of information encoded before the rest period^37–40^.

In contrast to multiple studies demonstrating that targeted memory reactivation during sleep benefits memory, less work has examined the impact of stimulus re-exposure during awake periods. Of the work that has examined awake re-exposure, mixed results have been reported: some studies found no significant impact of cueing^19,25,41–43^, with one study reporting a positive impact on reward-related memories^44^, and other work suggesting that awake cueing may impair or destabilize cued memory representations^23,45^. However, the majority of studies performed cueing during an attention-demanding working memory task^19,23,44^. With the exception of two related studies^25,41^, the content of awake periods when cueing is performed has not been carefully considered or manipulated. Thus, it is unclear whether the success of awake memory cueing depends on the behavioral state during cueing.

We hypothesized that, as opposed to a vigilant or externally-oriented awake state (e.g. an attention-demanding working memory task), a more quiescent awake state may be permissible to memory cueing and reactivation, as SWRs and reactivation tend to occur during pauses in ongoing behavior or periods of decreased task engagement^9,32–34^. Furthermore, as lower levels of acetylcholine are thought to promote hippocampal dynamics that facilitate reactivation and memory consolidation^30,33,46–48^, awake cueing may be most successful during time periods of reduced environmental novelty or a repetitive task, which should reduce levels of acetylcholine^49–51^. Here, we tested the hypothesis that awake targeted memory reactivation benefits memory when it is performed during a highly repetitive task, and that the degree of external attentional focus should be inversely related to cueing benefits.

To examine these questions, participants first underwent training to criterion to learn object-location associations (Fig. 1). After learning, an immediate memory test was administered to assess baseline object-location memory prior to cue re-exposure. Participants then performed a cover task (lexical decision task, LDT) during which recently learned objects were briefly visually re-exposed. The LDT was designed to minimize externally-oriented attention, yet ensured that participants were awake and viewing the computer screen. Lexical decisions were separated by long inter-trial intervals (ITIs) during which a repetitive arrow task was performed (Fig. 1). After the repetitive arrow task, a subset of recently learned object cues were briefly visually presented for 50ms and were immediately followed by the presentation of a word or pseudo-word stimulus displayed on a visual mask. We then examined the influence of object cueing during the LDT on delayed memory testing. We also asked whether reaction time (RT) on the arrow task, serving as an indirect measure of the degree of external attentional focus or vigilance, was related to the influence of cueing on later memory. Critically, given that active retrieval is known to benefit later memory^52^, we also assessed explicit recognition of cued items and asked whether cueing memory benefits were related to explicit cue memory. Lastly, we explored whether cueing benefits were related to initial learning.

**Figure 1.**
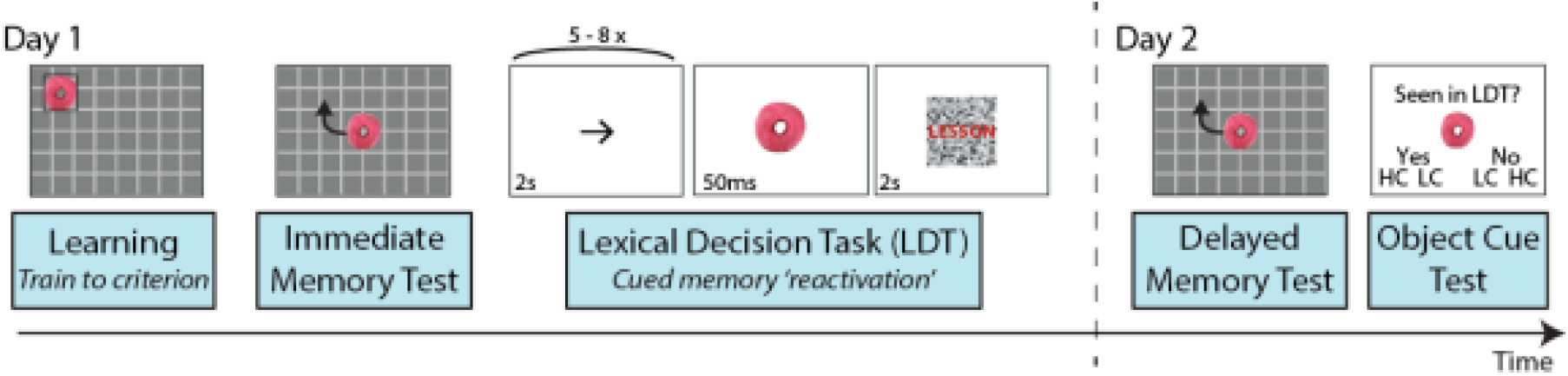
Experimental design. On Day 1, participants underwent training to criterion to learn 18 unique object-location associations (learning). Training consisted of consecutive rounds of learning and memory testing (with feedback). Associations were removed from training when each object was placed within 200 pixels of its assigned location during two consecutive memory tests. Learning was followed by an immediate memory test. During memory testing, each object was presented in the center of the grid and participants moved the object to its associated spatial location (no feedback was provided). A subset of objects was then briefly re-exposed or cued (for 50ms) during a repetitive foil task (lexical decision task, LDT) with each object re-exposed three times. Participants returned approximately 24 hours later for delayed memory testing, which consisted of two rounds of testing for each object-location association. Explicit knowledge or recognition of which objects were cued during the LDT was assessed during the object cue test, in which participants decided whether each object was seen or not seen during the LDT (high confidence, HC, or low confidence, LC).

## RESULTS

### Initial learning performance

All participants (N=23) learned 18 object-location associations via a training to criterion procedure. Training was completed in a total of 2 – 6 rounds of learning and test sessions (mean = 3.91 ± .23 rounds (standard error of the mean), median = 4 rounds), with an average of 2.55 ± .10 rounds for each association. During immediate memory testing, the average error or distance between studied and placed locations across all associations was approximately 77.3 ± 5.1 pixels, or 2.00 ± .13 cm. A small number of associations were poorly remembered during immediate memory testing. On average, almost one trial with relatively high spatial error (greater than two standard deviations from the mean distance across trials) was present for each participant (mean = .83 ± .14 trials; median = 1 trial; range = 0 – 2 trials). The average error of these trails was 264.3 ± 39.2 pixels (6.83 ± 1.01 cm). To avoid differentially biasing memory for cued or uncued associations, associations with poor initial memory were flagged and assigned as neither cued or uncued during subsequent comparisons (see Methods, Object cueing). Eight associations were then chosen to serve as cued and uncued associations based on equating memory during immediate testing.

### Delayed memory testing

To first understand the nature of how memory representations may change from initial learning to delayed memory testing (Fig. 1), we asked whether object placement during delayed testing was better predicted by each object’s studied location during *learning* or its placed location during *immediate memory* testing^53^. To do so, we compared the spatial distance or error between each object’s placed location during delayed memory testing to each of these positions (studied and remembered locations). When examining the mean error for each participant, a repeated measures ANOVA using within-subjects factors of memory assessment (distance computed relative to studied vs. remembered location) and testing round (first and seconds rounds of memory testing) showed a robust main effect of memory assessment (*F*(1,22) = 69.33, *p* = 3.02 × 10^−8^, ηp^2^ = .76). As shown in Figure 2a, the remembered location was a better predictor of placement during delayed memory testing for all participants. Interestingly, an interaction between memory assessment and testing round was also present (*F*(1,22) = 4.75, *p* = .04, ηp^2^ = .18), which was driven by an increase in spatial error from the first to second round of testing when memory was measured relative to each object’s studied location (*t*(22) = 3.06, *p* = .006, Cohen’s *d* = .64); however, spatial error remained stable across rounds of testing when measured relative to each object’s remembered location (*t*(22) = 1.52, *p* = .14, Cohen’s *d* = .32). This indicates that across repeated rounds of delayed memory testing, object placement drifted further from the studied location, but not the remembered location.

**Figure 2.**
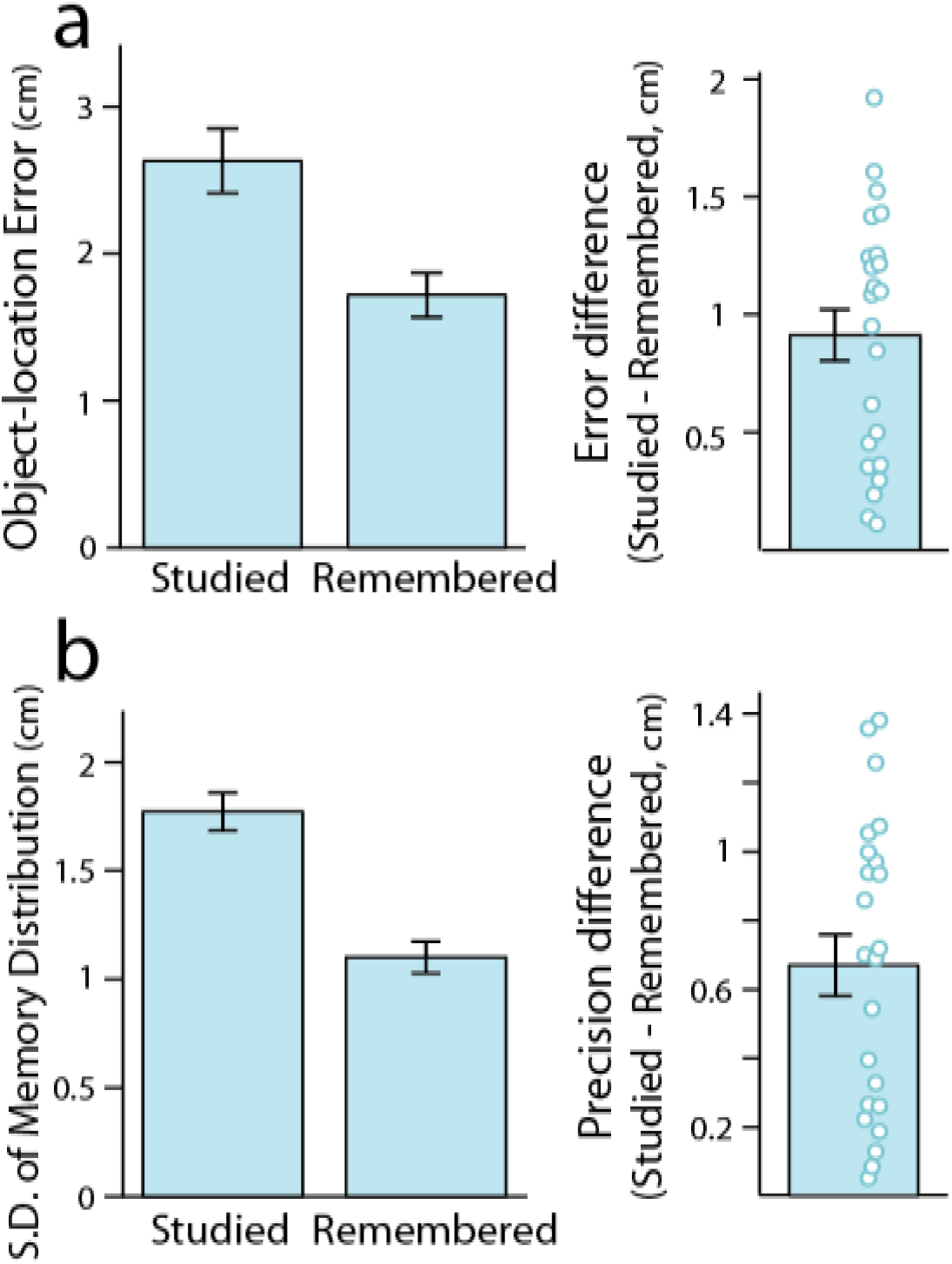
Delayed object placement was better predicted by remembered versus studied locations. (a) Average distance or error between object placement during delayed memory testing compared to its studied or remembered location. Left panel shows average across participants, right panel shows difference between studied relative to remembered locations, with individual data points corresponding to each participant. (b) Object-location memory precision estimated from fitting probabilistic mixture models during delayed testing (standard deviation of fitted Gaussian distribution) based on studied location or remembered locations. Lower standard deviations represent higher memory precision. As in (a), left panel shows average precision across participants and (b) shows difference in memory precision relative to studied versus remembered locations for individual participants. All error bars represent standard error of the mean across participants.

When considering individual trials, spatial error relative to placement at immediate memory testing was smaller than relative to its learned location for 73.6% of trials (71.7% for the first testing round and 75.4% for the second testing round). We also fit probabilistic models to model object placement as a mixture of guessing behavior and successful associative memory, separately considering memory relative to studied and remembered locations (see Methods, Statistical Analysis: Mixture Models). Greater memory precision was found when modeling delayed spatial placement relative to remembered versus studied locations (Fig. 2b; standard deviation of fitted Gaussian distribution for studied location = 1.77 ± .09 cm, remembered location = 1.1 ± .07 cm; *t*(22) = 7.53, *p* = 1.58 × 10^−7^, Cohen’s *d* = 1.57).

These results indicate that delayed object-location memory representations more closely matched memory expressed after training to criterion, rather than a veridical representation of the location studied during training. Since remembered compared to studied locations better predicted object placement across repeated trials of memory testing, it suggests that remembered locations reflect a more robust and resilient memory representation. Given these findings, we thus focused our analysis of the impact of cue re-exposure on subsequent memory by examining memory stability, or the change in delayed object placement relative to each object’s remembered location. However, as prior work has examined how cue re-exposure influences veridical memory, as a secondary goal, we also assessed the influence of cue re-exposure on later memory by measuring delayed object placement relative to each object’s studied location^19,44,54^.

### Influence of cueing on memory

Our primary question was whether brief visual re-exposure of object cues, or awake targeted memory reactivation, would enhance later object-location associative memory for cued compared to uncued associations. We first ensured that immediate memory measured before cueing was equated between cued and uncued associations (distance between remembered and studied locations for cued associations = 1.81 ± .10 cm, uncued associations = 1.81 ± .10 cm). Considering average delayed memory stability or graded spatial error during delayed memory testing (relative to object placement during immediate memory testing), a repeated measures ANOVA with factors of cueing and testing round showed a main effect of cueing (*F*(1,22) = 5.99, *p* = .023, ηp^2^ = .21) with no reliable cueing by testing round interaction (*F*(1,22) = 1.75, *p* = .20). As shown in Fig. 3a, a reduction in average spatial error or greater memory stability from immediate to delayed memory testing was found for cued relative to uncued associations. Similarly, a main effect of cueing on the probability of associative memory success derived from fitting mixture models was found (*F*(1,22) = 4.43, *p* = .047, ηp^2^ = .17), with a higher probability of memory success for cued (95.4 ± 1.2%) versus uncued associations (92.9 ± 1.7%; benefit of 2.5 ± 1.1%).

**Figure 3.**
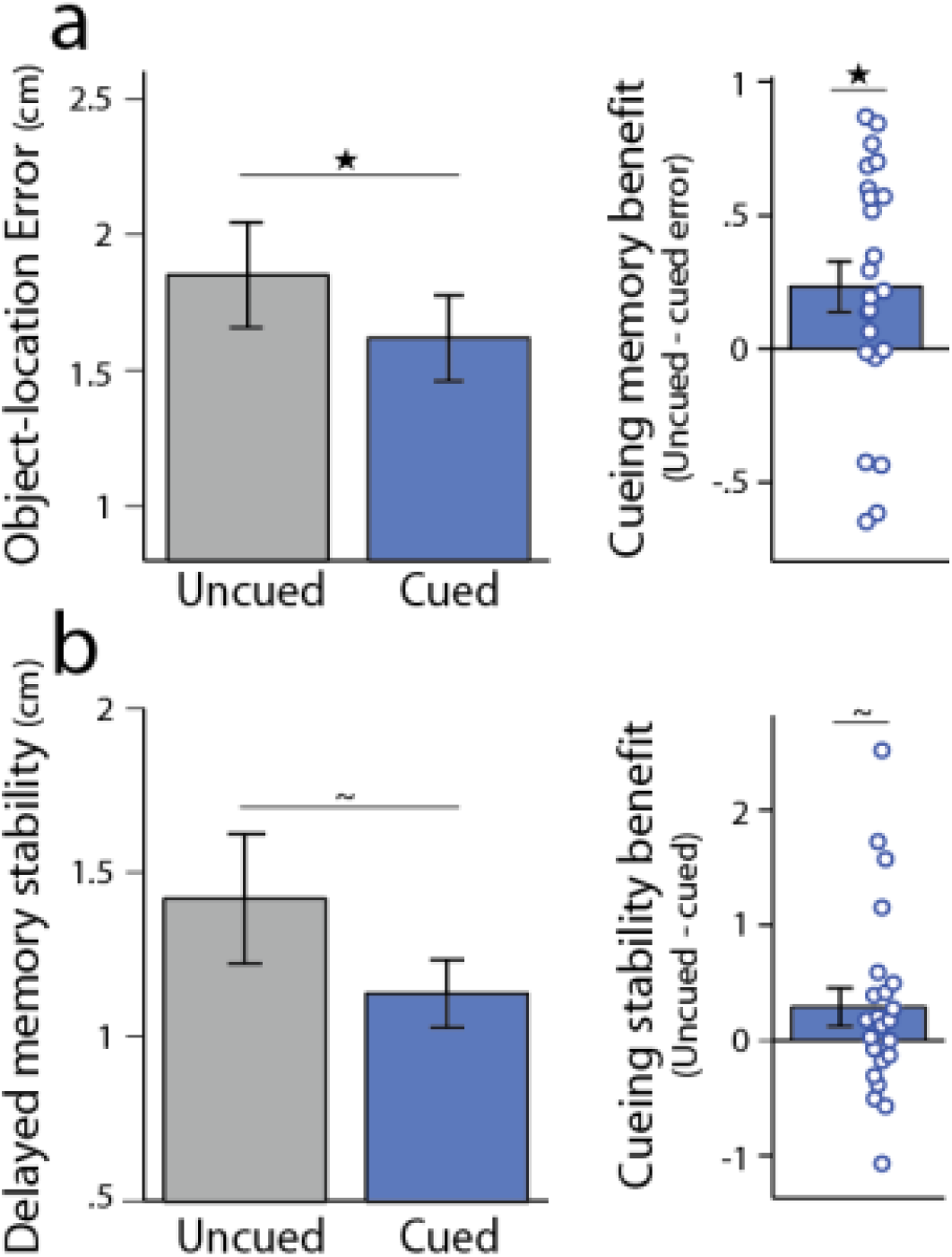
Object-location memory stability for cued versus uncued associations. (a) Average object-location error or memory stability (distance between object placement during delayed and immediate memory testing) for cued and uncued associations. (b) Average object-location memory stability across repeated trials during delayed testing for cued and uncued associations (distance between object placement across trials). For both (a) and (b), left panel shows average error or distance across participants and right panel shows cueing benefit (difference in memory for uncued minus cued associations), with each dot corresponding to the difference for each participant. 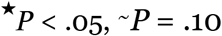

In addition to changes in memory stability from immediate to delayed memory testing, we also asked whether cueing influenced the stability of object-location memory across repeated blocks of testing on Day 2. A trend was found for greater stability or decreased distance between repeated trials during delayed memory testing for cued versus uncued associations (Fig. 3b; *t*(22) = 1.72, *p* = .10, Cohen’s *d* = .36). Together, these results indicate that cueing or targeted memory reactivation during the LDT enhanced the stability of object-location memory representations.

Since prior work has focused on examining how targeted memory reactivation influences memory for the presented locations during study, we also assessed the influence of cueing on delayed object-location memory considering each object’s studied location^19,44,54^, rather than its remembered location during immediate memory testing^53^. Although graded levels of object-location error relative to their studied locations were numerically reduced for cued relative to uncued associations (Supplementary Figure 1), no significant main effect of cueing (*F*(1,22) = 1.43, *p* = .24) or reliable cueing by testing round interaction (*F*(1,22) = 2.22, *p* = .15) was found using a repeated measures ANOVA with factors of cueing and testing round. However, a trend-level main effect of cueing was found on the probability of associative memory success derived from fitting mixture models (memory modeled relative to studied location; *F*(1,22) = 4.02, *p* = .057, ηp^2^ = .15), with a higher probability of memory success for cued (97.3 ± 1.2%) versus uncued associations (95.1 ± 1.3%; benefit of 2.2 ± 1.1%). These mixed findings suggest that the influence of awake cueing on veridical or studied object-location memory representations appear to be less robust or consistent across analysis approaches, as compared to the reliable influence of cueing observed across measures of object-location memory *stability* from immediate to delayed testing. Given these results and that object placement during immediate testing was a much better predictor of delayed object-location memory as compared to studied locations (see Delayed memory testing, above), our subsequent analyses focused on measuring the influence of cueing on object-location memory stability rather than veridical or studied object-location memory.

### Task engagement prior to re-exposure is related to the impact of cueing on memory

After establishing that brief awake re-exposure of objects can enhance the stability and expression of object-location memory, we next asked whether ongoing behavior at the time of object re-exposure predicted the impact of cueing on later memory. We hypothesized that when participants are in a less vigilant state, or have a greater internal versus external attentional focus, re-exposure would have a greater benefit on memory stability. To test this prediction, we assessed response time (RT) during the arrow task performed prior to each re-exposure event (Fig. 1). We then computed the average RT across the three re-exposure events for each cued object (and normalized RTs within participants, see Methods), resulting in a measure of vigilance or task-related external engagement prior to re-exposure events on an item-by-item basis (for each cued association). We predicted that longer RTs, possibly reflecting greater disengagement from the LDT, would be related to a greater cueing benefit, compared to associations that were re-exposed when participants were in a more vigilant or externally oriented state (when RTs were faster). Across all associations, the RT *prior to* re-exposure events was positively predictive of greater object-location memory stability for subsequently cued associations (Fig. 4, left panel; robust regression, *β* = 5.77 ± 1.70 (s.e.), *r* = .25, *t*(182) = 3.40, *p* = .00084; permutation test, *p* = .0012). Although robust regression was used to minimize the influence of potential outlier data points when assessing this relationship (as seen in Fig. 4), the correlation remained significant when excluding data points that were three standard deviations from the mean in either measure (*β* = 4.51 ± 2.19, *r* = .15, *t*(175) = 2.06, *p* = .04; permutation test, *p* = .02). This result suggests that awake cueing may be most beneficial for subsequent associative memory stability when re-exposure occurs during a less vigilant or externally attentive state.

**Figure 4.**
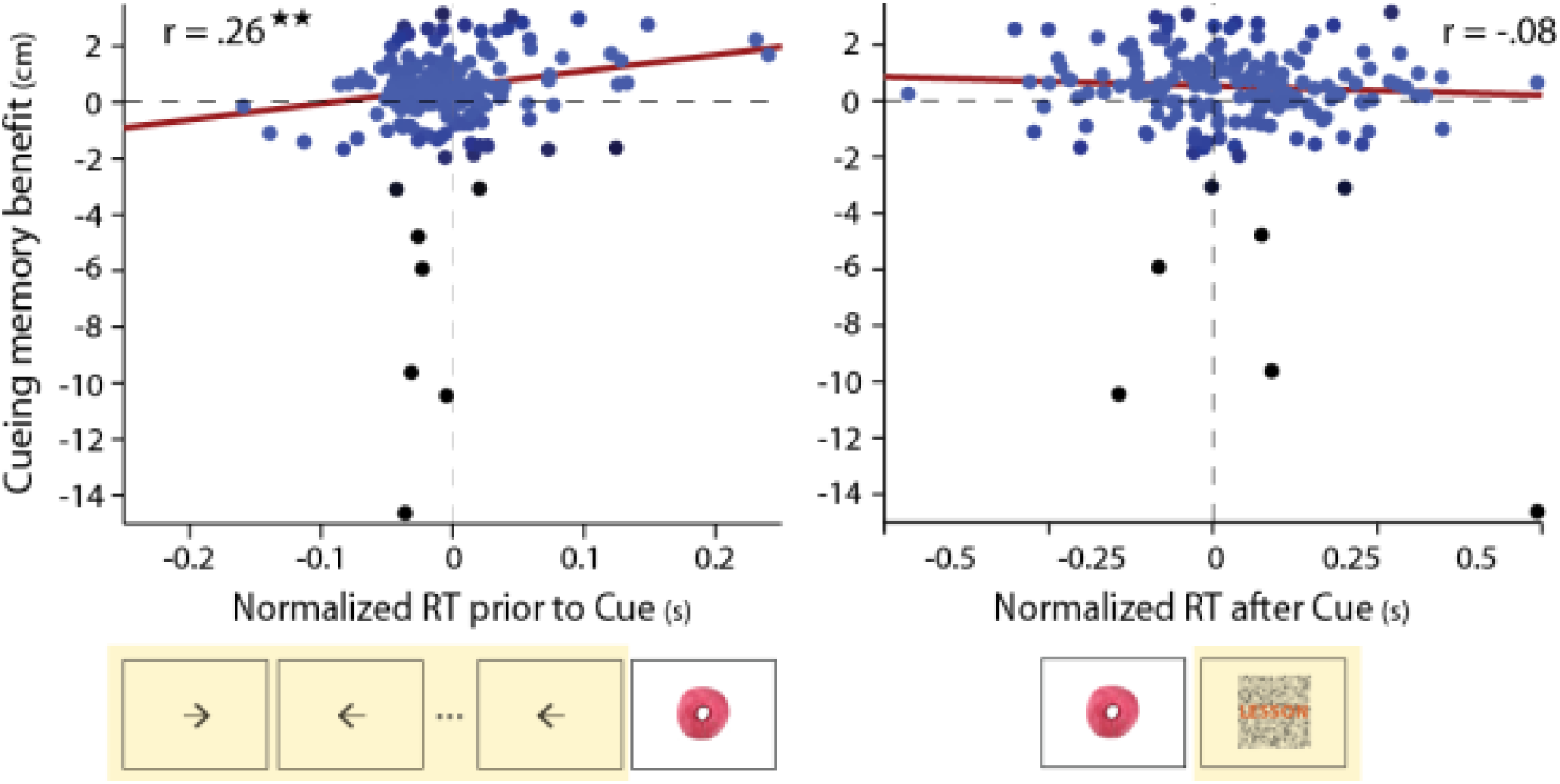
Cueing memory benefit as a function of ongoing behavior before and after object re-exposure. Data points correspond to the benefit in object-location memory stability for individual cued associations (mean uncued error minus error for each cued association, where error is the distance between object placement from immediate to delayed memory testing). The cueing benefit is shown as a function of mean normalized reaction time (RT) on the arrow task (performed during the ITI) prior to the re-exposure of each association (left panel, time period highlighted in yellow) and the mean normalized RT on the lexical decision task after the re-exposure of each association (right panel, time period highlighted in yellow). Robust regression was used to assess linear relationships between RT and cueing-related memory benefits. Each data point or association is colored based its weight or contribution to the regression model (black data points have a weight close to 0, while blue points have a weight of 1). 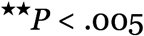

In addition to examining how a measure of behavioral state before re-exposure is related to the impact of cueing on memory stability, we next queried whether re-exposure impacted subsequent task performance, and whether this influence was related to benefits in memory stability for cued associations. Word decision trials that were preceded by object re-exposure showed a trend-level increase in reaction time compared to trials not preceded by re-exposure events (*t*(22) = 1.88, *p* = .073, Cohen’s *d* = .39; mean RT difference = 32.28 ± 17.14 ms). However, this tendency for re-exposure to influence subsequent task performance was not related to the impact of cueing on memory: a non-significant relationship was found between normalized RT on the subsequent word decision trial and cueing benefits for individual associations (Fig. 4, right panel; robust regression, *β* = −.65 ± .58 (s.e.), *r* = −.08, *t*(182) = −1.12, *p* = .26). These findings indicate that although re-exposure influenced ongoing processing during the LDT, this impact on behavior did not relate to cueing-related benefits in memory stability.

### Cueing benefits are related to initial learning

We next explored whether the influence of cueing on associative memory stability varied as a function of initial learning. Prior work has shown that associations with stronger or weaker initial memory strength may benefit differentially from post-encoding manipulations^22,54^ and that individual differences in the extent of initial learning are related to cueing benefits^54^. To test the latter idea, we examined two measures: object placement during immediate memory testing (relative to its studied location) and the average number of training blocks needed to reach learning criterion. We first assessed the relationship between these measures of learning and found that they were correlated (robust regression, *β* = .41 ± .10, *r* = .69, *t*(21) = 4.14, *p* = .00046). We thus derived a single factor score representing learning/initial object-location memory across these measures (using principal components analysis, see Methods). As shown in Fig. 5a, individual differences in object-location learning/initial memory were indeed related to cueing benefits (robust regression, *β* = .19 ± .06, *r* = .60, *t*(21) = 3.35, *p* = .003), with the worst learners showing the strongest benefits in memory stability from re-exposure. Similar results were obtained when this relationship was examined with each measure of learning separately: the cueing benefit was predicted by object-location error during the immediate memory test (*β* = .34 ± .15, *r* = .45, *t*(21) = 2.33, *p* = .03) and the average number of blocks to reach learning criterion (*β* = .50 ± .16, *r* = .59, *t*(21) = 3.22, *p* = .0041). These findings indicate that memory benefits associated with awake re-exposure were strongest for individuals with the worst learning or initial memory performance.

**Figure 5.**
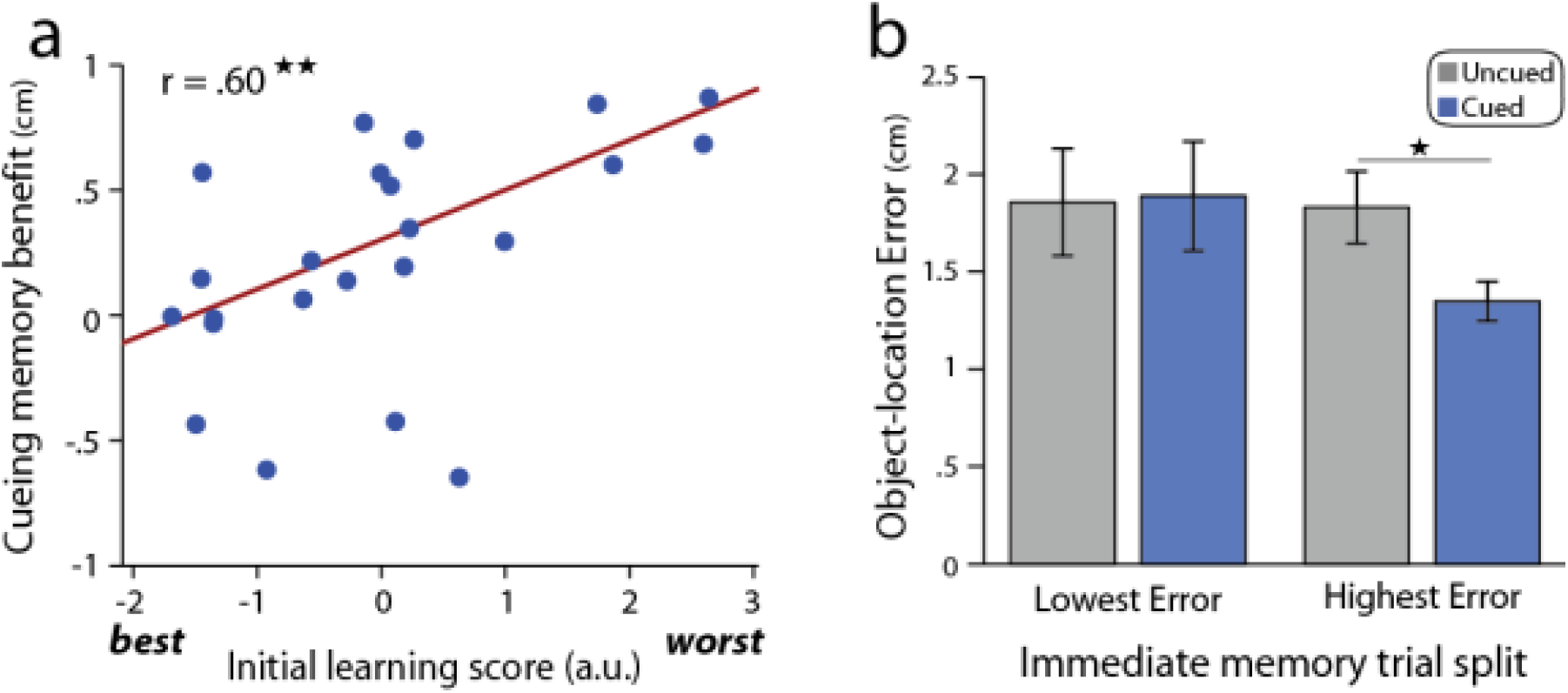
Cueing memory benefits are related to initial learning or memory. (a) Individual differences in the cueing memory benefit (mean uncued minus cued error or distance between object placement from immediate to delayed memory testing) are shown as a function of initial learning/memory performance. The initial learning score represents a single factor score or the first principal component derived across learning measures (mean spatial error during immediate memory testing and the average number of rounds to reach learning criterion). Lower scores indicate better memory and faster learning, while higher scores indicate worse memory and slower learning. (b) Cueing memory benefits as a function of item-level differences in initial memory. A split half analysis was performed on cued and uncued associations for each participant. Cueing did not reliably influence memory stability (distance between object placement from immediate to delayed testing) for associations with the lowest error or best memory during initial memory testing (left bars), while a benefit in memory stability was found for cued versus uncued associations with the highest or worst memory during initial testing (right bars).

In addition to examining whether individual differences in initial memory were related to cueing benefits, we also assessed whether similar effects were seen for individual associations within participants, as prior work has found that relatively weaker associations benefit most from cueing^22,54^. To do so, we performed a split-half analysis on cued and uncued associations within each participant based on spatial error during immediate memory testing (Fig. 5b). A repeated measures ANOVA on delayed memory stability with within-subjects factors of initial memory (low vs. high error), cueing, and testing round showed a trend for an interaction between cueing and initial memory (*F*(1,22) = 3.44, *p* = .077, ηp^2^ = .14) as well as a main effect of cueing, consistent with our prior analysis (*F*(1,22) = 5.80, *p* = .025, ηp^2^ = .21). As shown in Fig. 5b, cueing benefited memory for high error trials (*t*(22) = 2.71, *p* = .013, Cohen’s *d* = .56), but did not reliably impact memory for low error trials (*t*(22) = 0.21, *p* = .83, Cohen’s *d* = .05). These results indicate that cueing most reliably benefited associative memory stability for the weakest associations, whereas the most strongly learned associations were not reliably influenced by re-exposure.

### Cueing benefits are not driven by explicit cue knowledge

Lastly, an important question to consider is how re-exposure during an awake time period may impact later associative memory: through explicit recognition and re-encoding of the cue stimulus, which may trigger overt retrieval of object-location representations, or whether re-exposed cues do not result in explicit retrieval, but instead serve to subtly bias brain activity and potentially promote the reactivation of cued associations. First, our prior result that the influence of cueing on subsequent reaction time was not predictive of the memory benefit associated with cueing (Fig. 4) speaks against the notion that cues triggered effortful retrieval, and that any effortful retrieval may have resulted in enhanced memory stability for cued associations.

To assess participant’s awareness of the cueing manipulation, we administered a questionnaire after delayed memory testing. The majority of participants (78% or 18/23) reported noticing that objects seen during learning were presented again during the LDT. About half of the participants (52% or 12/23) were able to recall at least one object that was re-exposed during the LDT. Thus, it is clear that many participants were explicitly aware of the brief re-exposure manipulation, leaving open the possibility that explicit knowledge of cued items was responsible for cueing-related associative memory benefits.

In order to assess knowledge of which items were re-exposed during the LDT, participants performed a forced-choice task in which each object was rated as either seen or not seen during the LDT. This allowed us to examine relationships between explicit cue knowledge and cueing benefits, both across participants and at the level of individual associations. Across all participants, reliable discrimination between cued and uncued objects was evident (Fig. 6a, cueing by rating interaction, *F*(1,22) = 18.42, *p* = .0003, ηp^2^ = .46). Cued objects were more likely to be rated as high confident cued than uncued objects (*t*(22) = 4.22, *p* = .00035, Cohen’s *d* = .88) and uncued objects were more likely to be rated as not seen during the LDT compared to cued objects (low confident responses, *t*(22) = 2.51, *p* = .02, Cohen’s *d* = .52; high confident responses, *t*(22) = 2.46, *p* = .022, Cohen’s *d* = .51). However, despite this reliable discrimination at the group level, considerable variability in successful cue discrimination was present across participants (Fig. 6b).

**Figure 6.**
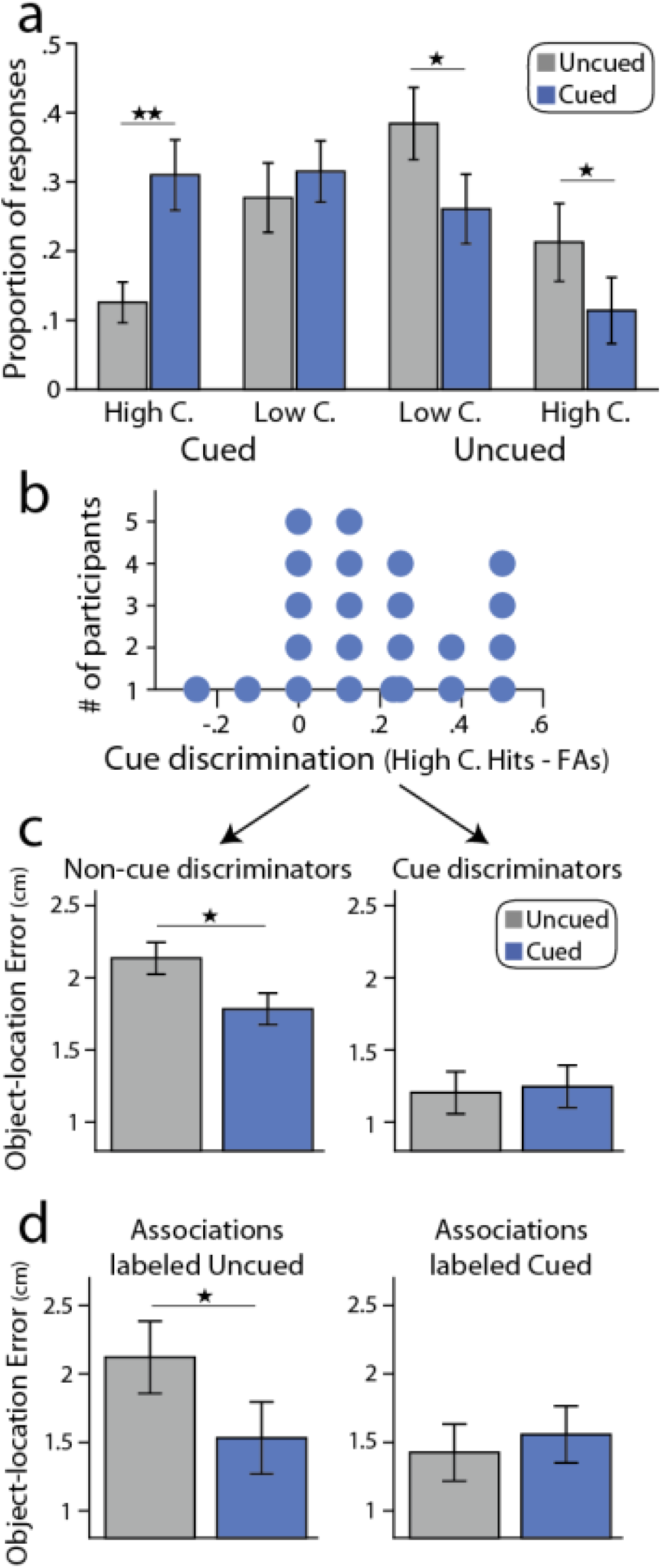
Relationships between cueing memory benefits and explicit cue knowledge. (a) Distribution of responses on cue discrimination test for cued and uncued objects. On average, participants were more likely to rate cued objects as high confident cued (High C., left bars) and were more likely to rate uncued items as high and low confident uncued (right bars). (b) Histogram of cue discrimination performance across participants (high confident hits minus false alarms). (c) Object-location memory stability (distance between object placement from immediate to delayed testing) shown separately for participants without reliable or above-chance cue discrimination (Non-cue discriminators, left panel) versus participants with reliable cue discrimination (Cue discriminators, right panel). A marginal difference in cueing memory benefits was found between non-cue versus cue discriminators (see Results). (d) Object-location memory stability shown for associations labeled as uncued (left panel) and associations labeled as cued (right panel).

Given the variability in cue discrimination across participants, we asked whether re-exposure benefits were reliable when restricting our analysis to participants that did not show above chance cue discrimination. We categorized participants based on whether or not they showed reliable above chance cue discrimination using a chi squared test. As shown in Fig. 6c, participants without reliable cue discrimination showed enhanced object-location memory stability for cued versus uncued associations (*t*(15) = 3.2, *p* = .006, Cohen’s *d* = .67), whereas re-exposure did not reliably influence object-location memory stability for participants with above chance cue discrimination (*t*(6) = −0.29, *p* = .78, Cohen’s *d* = .06). A nearly significant trend for differential cueing benefits across participants with versus without reliable cue discrimination was found (*t*(21) = 2.05, *p* = .053, Cohen’s *d* = .95). These results importantly indicate that the cueing memory benefit was not driven by participants with explicit recognition of which objects were re-exposed. Rather, cueing benefits were most reliable for participants without the ability to discriminate cued from uncued objects. However, as is evident in Fig. 6c, it is important to note that differences in overall memory stability were present across participants that were able to reliably discriminate cues versus those without reliable discrimination (comparison of average object-location memory stability across cued and uncued associations, *t*(21) = 2.14, *p* = .044). Thus, it is unclear if the difference in cueing benefits across cue versus non-cue discriminators may be indirectly related to differences in overall memory stability or task performance across these groups. Nonetheless, our results speak against the interpretation that cueing benefits were particularly driven by participants that reliably discriminated between cued and uncued objects.

To further probe whether cueing memory benefits were driven by explicit recognition of cued items and thus potentially explicit retrieval during cueing, we examined the influence of cue recognition on memory benefits at the level of individual associations. To assess whether correctly identified cued objects were driving the cueing benefit, we excluded associations in which the object was labeled as cued. As shown in Fig. 6d, a reliable cueing benefit was present when only associations labeled as uncued were examined (*t*(20) = 2.14, *p* = .045, Cohen’s *d* = .45). No difference in memory was found for associations labeled as cued (*t*(18) = 0.42, *p* = .68, Cohen’s *d* = .09), although no reliable difference in the cueing benefit for associations labeled as cued versus uncued was found (*t*(16) = 0.97, *p* = .35, Cohen’s *d* = .20). Unlike the comparison across participants as a function of reliable cue discrimination (Fig. 6c), this item-level analysis was not confounded by overall differences in delayed memory; no reliable difference in delayed object-location memory was present across associations labeled as cued versus uncued (*t*(20) = 1.18, *p* = .25). These findings critically demonstrate that benefits of re-exposure on memory stability were not explicitly driven by overt recognition of cued items, as reliable memory benefits remained when associations labeled as cued were excluded.

## DISCUSSION

Much recent work has examined the impact of stimulus re-exposure during sleep on subsequent memory, demonstrating that targeted memory cueing benefits later memory and providing compelling evidence that off-line reactivation may be an important mechanism supporting memory^19,21,44^. In contrast, it is less clear from prior work whether similar cueing during the awake state benefits memory. Here, we demonstrate that brief (50ms) visual cueing while participants are awake and performing a repetitive task can benefit associative memory stability. Importantly, explicit knowledge of cued items did not drive cueing benefits, suggesting that the memory benefit was not driven by active retrieval or cue-triggered rehearsal processes. These findings extend our understanding of the ways in which targeted memory reactivation can benefit memory, and in doing so, provide novel evidence that reactivation during awake post-encoding time periods contributes to memory stabilization.

An important issue to consider is whether any influence of awake cueing on later memory was driven by explicit retrieval and rehearsal processes during cueing, as active retrieval is known to benefit memory^52^. Our participants were, on average, able to discriminate between cued and uncued items, in contrast to prior work performing cueing during sleep^19,22,54^. However, we think that it is unlikely that our results were driven by cue-triggered rehearsal or explicit recognition of cues for several reasons. First, cueing-related memory benefits were greater in participants that could not reliably discriminate cued from uncued items compared to participants with above-chance recognition of cues, suggesting an inverse relationship between explicit cue recognition and cueing-related memory benefits (however, we note that this comparison was confounded with overall differences in delayed memory across these participants, see Fig. 6c). If cueing elicited active retrieval of object-location associations, participants with the highest recognition of cued items should show the greatest cueing memory benefit, but the opposite pattern of results was observed. Secondly, cueing benefits remained when restricting our analysis to associations in which the object was labeled as not cued. Moreover, only two participants reported that the re-exposed objects caused them to explicitly think about their associated spatial locations from prior learning. Finally, the immediate influence of cueing on subsequent behavior during the LDT was not related to cueing memory benefits. It thus appears unlikely that cue-triggered retrieval was responsible for driving cueing-related memory benefits. This evidence against a relationship between explicit cue-triggered retrieval and the benefit of awake cueing on memory parallels the result that rehearsal during post-encoding rest periods is not responsible for driving the benefit of awake rest periods on subsequent memory^38^ and is consistent the demonstration that conscious access to cued memories is not necessary for their subsequent modification after cueing^55^.

Most work examining the influence of cueing during awake periods has performed cueing during an externally-oriented, attention-demanding task, with the goal of minimizing explicit recall and/or rehearsal of cued associations^19,44^. However, it is unclear whether cueing during an externally oriented attention-demanding task can influence sensory representations and potentially bias ongoing or induce hippocampal reactivation events. Here, we sought to bias ongoing brain activity by presenting brief (50ms) re-exposures during a task that was designed to be repetitive and promote an internally versus externally oriented attentional state. We found that greater cueing benefits were associated with longer response times on the task performed prior to cueing, consistent with the hypothesis that a state of reduced external engagement or vigilance may be most beneficial for memory cueing and reactivation. These findings are also congruent with the notion that reduced cholinergic tone associated with reduced external attention demands and novelty^49–51^ may promote a brain state that is permissible to consolidation-related mechanisms^30,33,46–48^. However, it is important to note that response times serve as an indirect measure of vigilance or a proxy for an internally versus externally orientated behavioral state. Longer response times may also reflect other factors, such as the degree of external distraction, interference from the prior lexical decision, task difficulty, or overall fatigue. Thus, future work is needed to more directly test the hypothesis that internal versus external attentional focus, or levels of vigilance, are related to the influence of cueing on memory.

In this framework, it is unclear why prior studies that performed cueing during task-free awake periods did not influence memory^25,41^. One of several differences between our study and the cueing performed by the studies of Schreiner & Rasch may have contributed to diverging findings. We presented brief (50ms) visual re-exposures over ~25 minutes, with a rate of approximately one cue per minute, while Schreiner & Rasch performed cueing over 90 minutes with re-exposures once approximately every three seconds. Although their cueing schedule was successful during sleep, it is possible that this rate of awake cueing was too rapid to influence reactivation, or occurred so frequently that cues were actively suppressed. Another possibility is that cueing during a true awake rest period, when a participant can fully attend to cues, may not be optimal for inducing reactivation. Rather, this state may promote explicit retrieval, which may not benefit memory if retrieval is not successful or if non-task-relevant information is retrieved. An additional difference is that we used a graded measure of associative memory stability, while Schreiner & Rasch assessed veridical measures of cued recall and recognition; it is possible that an influence of awake cueing would have been revealed using a more fine-grained memory measure or an assessment of memory stability. Finally, as Schreiner & Rasch tested memory shortly after cueing, it is possible to speculate that the influence of awake cueing on memory is stronger with longer delays. In addition to influencing reactivation during the cueing period itself, it is possible that awake cueing may serve to ‘tag’ or bias specific associations for subsequent reactivation and consolidation processes, such as those taking place during sleep after cueing and before delayed memory testing^56,57^. Prior work has shown that post-encoding manipulations can serve to preferentially enhance memory for particular subsets of recent experience, depending on their instructed relevance for future behavior^58–60^ or indirect associations with salient outcomes, such as aversive shocks or reward^61,62^. Thus, it is possible that brief visual re-exposures performed in the current study acted as salience signals to ‘tag’ cued stimuli or associations for later consolidation during future periods of sleep and/or wakefulness. As we only tested memory 24 hours after the cueing procedure, it would be interesting to examine this question in future work by testing memory at various delays after cueing.

It is also important to consider other differences between our approach and prior work that may have resulted in the observed influence of cueing. Before examining the influence of cueing on later memory, we first sought to understand the nature of delayed object-location representations in memory, given the measure of cued object-location recall being assessed. We were motivated by prior work^53^ to ask whether delayed object placement more closely matched each object’s studied location or its remembered location during immediate testing. Replicating this prior work^53^, we found that delayed object-location memory more closely matched memory expressed during immediate testing as compared to studied object locations. This is an important finding since, theoretically, cueing serves to reactivate an internal memory representation and if there is a substantial departure in initial memory from the actual veridical studied location, cueing can only influence the internal representation in memory. Consistent with this idea, we found a reliable influence of cueing on measures of memory stability (changes in object placement from immediate to delayed testing) and a more mixed influence of cueing when delayed memory was measured relative to studied object locations rather than remembered locations expressed during immediate testing. It is possible to speculate that the mixed effects of awake cueing seen across prior studies may be related to the use of studied or veridical memory measures rather than an assessment of memory stability. However, it is important to note that this discrepancy across memory measures in our study may also be related to the relatively liberal nature of the learning criterion used in during training (200 pixels, while prior studies have used a criterion of 150 pixels^44,54^). With the use of a more stringent learning criterion, internal memory representations should more closely match studied information and such large discrepancies in delayed memory measured relative to studied versus remembered locations (see Fig. 2) may not be present. Nonetheless, in order to fully understand the influence of cueing on later memory, future work should consider assessing measures of both veridical memory as well as memory stability from before to after cueing.

Another difference between previous work using targeted memory reactivation and the current study is that we briefly visually re-exposed participants to stimuli from prior learning, whereas prior investigations have primarily used either olfactory^42,45^ or auditory^19,22,25,41,44^ cues. Given that past studies have focused on cueing performed during sleep, when visual stimulation would not be practical, the use of auditory and olfactory stimuli which can modulate brain activity without inducing awakenings is warranted^20^. It is possible that brief visual cues may be better suited for influencing reactivation or for tagging associations for subsequent consolidation than cueing in other sensory modalities during wakefulness, which could potentially explain the lack of cueing benefits observed using auditory cues in prior work^25,41^. Regardless, our findings suggest that brief visual presentation may be an effective way to perform targeted memory reactivation during awake periods.

Our finding that cueing during the awake state can enhance later memory for object-location associations parallels and was motivated by prior work demonstrating that similar manipulations during NREM sleep improve memory for cued associations^19,21–25,42^. This similar finding across NREM sleep and awake periods designed to minimize externally-oriented attention suggests that cued memory reactivation may behave in a comparable manner across these brain states. Indeed, hippocampal reactivation and SWR activity, which are thought to be key mechanisms supporting memory stabilization, occur during both NREM sleep and quiescent awake periods, particularly during pauses in ongoing behavior or periods of decreased task engagement^8–10,32–34^. It has been proposed that mechanisms supporting memory consolidation may indeed occur during a brain state characterized by relatively low levels of acetylcholine^46,47^ when the hippocampus is not engaged in encoding novel information^7,30,31^. It is thus possible to view periods of quiescent wakefulness and NREM sleep as potentially belonging to a similar brain state, with NREM sleep representing a special case of reduced environmental novelty and low levels of acetylcholine^30^. Prior work directly comparing the influence of targeted memory reactivation across NREM sleep and wakefulness has performed awake cueing under a variety of conditions, typically during an attention-demanding working memory task^19,23,44^. These direct comparisons of awake and NREM sleep cueing have yielded mixed results; in most cases, no overall influence of awake cueing on later memory was found^25,41–43^. However, one study found that awake cueing more focally enhanced memory relative to cueing during sleep^44^, with other studies suggesting that awake cueing may destabilize representations in contrast with a stabilizing influence of cueing during sleep^23,45^. Lastly, another study found a numerical but non-significant influence of awake cueing on memory, in contrast to a significant influence of cueing during sleep^19^. Our findings suggest that it may be informative to first understand the time periods during wakefulness that are most reliably influenced by exogenous memory cueing, or are most permissible to endogenous reactivation, before assessing the similarities and differences that reactivation may serve across sleep and awake periods.

We also explored whether the success of cueing varies based on levels of initial associative learning. We found that both participants and associations with the poorest initial learning benefited the most from stimulus re-exposure. Differential cueing benefits at the level of individual associations within each participant complement recent work showing that cueing during sleep most strongly benefits the poorest learned associations within each participant^22,54^. Since prior work has shown that the strength of endogenous post-encoding reactivation varies with the strength of initial representations present during learning or encoding^13,63–65^, it is possible to speculate that both awake and sleep cueing does not benefit the strongest memories as they may already undergo sufficient endogenous reactivation. Thus, cueing of comparatively weaker representations may serve to preferentially promote or equalize their reactivation relative to stronger representations. Our data show congruent relationships between initial learning and cueing benefits both at the level of individual associations as well as across participants. In particular, we found that participants with the poorest initial learning benefitted the most from awake cueing. This finding stands in contrast to recent work showing that participants with the highest initial memory performance most strongly benefit from cueing during NREM sleep^54^. It is currently unclear whether these distinct results across studies at the level of individual differences across participants represent a true difference between awake versus sleep cueing or were driven by procedural differences across studies (learning criterion, memory measures, and delays between learning, cueing, and memory testing varied between studies). For example, our learning criterion (200 pixels) was more liberal than in prior work (e.g. 150 pixels^54^) which may allow cueing to more strongly benefit participants or associations with overall weaker levels of learning. Future work is needed to understand the factors driving differences between our studies. Regarding our finding that cueing benefits participants with the weakest initial learning, it is possible that participants with the strongest initial encoding already undergo sufficient reactivation and consolidation-related processes during post-encoding time periods, and cueing thus does not additionally benefit memory for these participants. This idea is consistent with work showing that both endogenous and exogenous (cued) post-encoding reactivation is greatest in response to higher learning demands^27,66–68^.

Lastly, it is interesting to consider potential parallels between the approach of targeted memory reactivation and techniques used for studying memory reconsolidation or updating. As in the current study, targeted memory reactivation seeks to induce neural reactivation of learned representations via the presentation of a cue associated with a learning context (e.g. an odor) or individual associations (typically sounds). The goal of this manipulation is to mimic or bias the endogenous memory reactivation that occurs during offline brain states via the presentation of an exogenous cue, and putative ensuing reactivation is thought to strengthen cued representations^20,26^. Memory reconsolidation or updating is similarly examined by presenting reminder cues (during awake periods), with the goal of returning memories to a labile state in which they may undergo modification^69^. To test for modification, cues are typically combined with physiological interventions to influence particular neural processes, or are paired with novel or interfering information. Memory is later probed to reveal an influence of modification or the incorporation of novel information into cued memories (for example see^70,71^). It is possible that the procedure used in the current study may have also resulted in the updating of cued object representations as well as strengthening of object-location associations. However, as we did not explicitly link the presentation of object cues with specific information and did not probe memory in a manner to reveal updating, it is unclear if our brief (50ms) cueing procedure would be effective for both memory strengthening and updating. Prior work has shown that both updating and strengthening of item representations can co-occur as a function of direct item reactivation (for longer reactivation durations of 5s^72^), so it is plausible that our procedure could lead to both strengthening and updating. However, other work suggests that the presentation of interfering information after awake cueing may compete with previously learned information^45^, suggesting a possible trade-off between strengthening of initial memory representations and updating (see also^24^). As memory updating may be driven by the presence of prediction error, or a mismatch between cued and subsequently presented information^73,74^, it is likely that distinct mechanisms may mediate memory strengthening versus updating. Future work is needed to directly explore relationships between memory strengthening and updating as a result of awake cueing procedures.

In conclusion, this study demonstrates that stimulus re-exposure during a post-encoding awake period can enhance the stability of memory representations. Importantly, our data indicate that it is unlikely that cue-triggered retrieval drove the influence of cueing on later associative memory. Our finding that the magnitude of external task engagement prior to cueing, as measured by response time, was inversely related to memory benefits is consistent with the notion that awake cueing may be most effective and reactivation may be most permissible during a more internally versus externally oriented brain state. This finding mirrors the occurrence of awake reactivation and SWRs during pauses in behavior^10,32,34^ and the notion that reduced cholinergic tone promotes hippocampal dynamics that support memory reactivation and consolidation^46,47^. However, future work is needed to explore more direct links between vigilance, internal versus externally oriented brain states, and memory reactivation. This work adds to a growing body of literature that post-encoding awake time periods may play an important role supporting reactivation and long-term memory^15,30,35,37,38^.

## METHODS

### Participants

Twenty-five native English speakers with normal or corrected-to-normal vision participated in the study. Two participants were excluded due to a large number of missed responses on the lexical decision task (number of missed responses > two standard deviations above the mean of all participants). The average age of the remaining 23 participants was 20.7 years (range: 18-34) and included 8 males and 15 females. Informed consent was obtained from all participants in a manner approved by the University Committee on Activities Involving Human Subjects (UCAIHS) at New York University. All experiments were carried out according to guidelines approved by UCAIHS.

### Procedure

Participants were informed of the general study procedures. On Day 1, participants learned object-location associations via a training to criterion procedure, followed by an immediate memory test, and the lexical decision task which served as a cover task for object cueing (Fig. 1). On Day 2, participants performed object-location memory testing, followed by an object cue recognition test, and an additional object recognition test (not presented here). Participants were not informed on Day 1 that their memory for object-location associations would be tested on Day 2.

### Stimuli

Object stimuli consisted of 18 unique, everyday objects on white backgrounds and were sized 250 × 250 pixels. Words used in the lexical decision task were chosen from the MRC Linguistic database and consisted of adjectives of 5 – 8 characters in length.

### Object-location learning

During repeated learning and test blocks, participants underwent training to criterion to learn 18 unique object-location associations. Each object was randomly assigned to a unique spatial location on the grid (referred to as its studied location). During learning trials, each object was shown in the center of the grid for 3 seconds. The object then moved to its associated spatial location over the course of 1 second. A red frame appeared around the object after it reached its associated location and remained in this location for a minimum of 2 seconds. Participants then clicked on the object to advance to the next trial. After an inter-trial-interval (ITI) of 2s, the next object appeared.

Each learning block was followed by a test block during training to criterion. During each trial in the test session, an object was presented in the center of the grid and participants were instructed to position the object in its studied location. After initially positioning the object, the mouse cursor appeared and participants clicked on the object to finalize its location or clicked and dragged the object to reposition it. Feedback was then provided and the object was displayed in its studied location. If the object was placed within 200 pixels of its studied location, a green frame appeared around the object to indicate that the position was approximately correct. If the object was positioned more than 200 pixels from its studied location, a blue frame appeared around the object and a black line was displayed between the studied and chosen locations. A 2s ITI occurred before the start of the next trial.

Training to criterion (rounds of learning and test blocks) continued until all objects were placed within 200 pixels of their assigned locations across two successive rounds of testing. Individual objects were removed from subsequent learning and test blocks after training criterion was reached.

### Object-location memory test

Memory for object-location associations was assessed on Day 1 (immediate memory test) and Day 2 (delayed memory test). Memory tests were identical to the test blocks during training to criterion, except that no feedback was provided. Specifically, each of the 18 objects was presented in the center of the grid, and participants were instructed to position each object in its associated location on the grid. The delayed memory test on Day 2 consisted of two rounds of testing.

### Object cueing

Like prior work^19^, we matched initial memory for object-location associations selected to be cued versus those that served as uncued or control associations. Out of the 18 object-location associations, eight were assigned to be cued, eight were assigned as uncued, and two were flagged to be neither cued or uncued (omitted from comparisons of cued and uncued associations). We excluded two associations from serving as cued or uncued since in pilot testing we often observed that one or two associations had very high spatial error during immediate memory testing, and we preferred to not include these associations as either cued or uncued to avoid biasing either group of associations. To assign associations to these categories, we first examined whether any objects had excessively high spatial error during immediate memory testing (if spatial error was two standard deviations above the mean across all trials). Any such associations were omitted, serving as neither cued or uncued associations. If this number of high spatial error associations was less than two, we randomly choose additional associations to serve as neither cued or uncued. We then assigned eight objects to be cued and eight other objects to be uncued based on their error during the immediate memory test. To achieve this, we tested combinations of cued and uncued assignments, and chose the assignment that minimized the difference in mean spatial error across cued and uncued associations during immediate memory testing.

### Lexical Decision Task

Objects were cued or briefly re-exposed after learning during an unrelated Lexical Decision Task (LDT; Fig. 1). This task was designed to be repetitive and promote an internally-oriented state, yet require that participants view the screen in order to visually cue object stimuli. The LDT lasted for approximately 25 minutes and consisted of 96 trials (word or pseudo-word presentations). Each trial of the LDT consisted of the presentation of red capital letters on top of a visually scrambled stimulus for 2s. Participants were instructed to indicate whether the letters were a word or non-word by pressing ‘w’ or ‘p’, respectively. The ITI was intentionally long and repetitive, lasting for either 10, 12, 14, or 16 seconds, and consisted of the presentation of a series of arrows presented on the screen. Each arrow pointed either to the left or right and was presented for 2s. A total of 5, 6, 7, or 8 arrows were presented in each ITI. Participants were instructed to press ‘w’ if the arrow pointed to the left or ‘p’ if the arrow pointed to the right. Long ITIs were used in order to promote a non-vigilant state and variable ITI durations prevented an expectation of the onset of lexical decision and the occurrence of object re-exposures.

On one-quarter of trials (24/96), objects were briefly re-exposed before the lexical decision trial (Fig. 1) for 50ms (three refreshes on a monitor with a 60 Hz refresh rate). Objects were visually masked as the words that immediately followed object re-exposures were presented on scrambled images of equal size of the objects. Each of the eight cued objects was re-exposed three times throughout the LDT. Object re-exposures occurred in a pseudo-random fashion as a function of task parameters (they were evenly distributed across the four ITI durations and occurred equally often before word and pseudo-word stimuli). Each of the three re-exposures for each object occurred pseudo-randomly over time throughout the LDT (one re-exposure for each one-third of task trials).

### Questionnaire

Before informing participants of the object re-exposure manipulation, we queried participant’s awareness of the manipulation using a questionnaire. The questionnaire contained several questions regarding participants level of engagement and thought content during the LDT and was administered after our main dependent measure of interest was collected (after delayed memory testing). We determined whether participants were aware of object re-exposures from responses to the following questions: “Did you notice a flash before some of the words were presented? If so, please describe what you saw and how this affected your experience?” and, for participants that noticed a flash before some words were presented, they answered “Did you notice anything in particular about the content of the flash(es)? Could you tell if any kind of visual information was displayed in the flash(es)?”.

### Object cue discrimination test

To assess explicit knowledge of which objects were cued during the LDT, participants rated each of the 18 objects as being cued or uncued during the LDT (whether or not each object was or was not seen during the LDT). The cue discrimination test occurred after delayed memory testing and completion of the questionnaire (Fig. 1) to ensure that our main dependent measure of interested (object-location delayed memory) was not biased by these decisions. Prior to the start of the discrimination test, we first informed participants that some of the objects seen during learning were briefly presented during the LDT (if they were not already aware of this manipulation, based on their responses to the questionnaire). During the discrimination test, each object was presented in the center of the screen, and participants rated each object as: high confident seen, low confident seen, low confident not seen, and high confident not seen (Fig. 1). Participants were instructed to take as much time as they needed in order to make the most accurate decision (no response time limit was given).

### Statistical Analysis: Graded object-location memory

We examined object-location memory during delayed memory testing to assess the impact of cue re-exposure on later memory. To first understand the nature of memory representations during delayed testing, we asked whether object placement was more similar to each object’s studied location (seen during training to criterion) or each object’s placement during immediate memory testing. This issue is relevant for our main question, as the extent to which object re-exposure serves to reactivate representations in memory, it may be most appropriate to measure delayed object placement relative the immediate memory test (as a proxy for its representation in memory), rather than the location studied during learning^53^. We thus computed the distance between each object’s placed location during delayed memory testing relative to its studied location as well as its remembered location (placement during the immediate memory test). Delayed object-location memory was then analyzed using a repeated measures ANOVA with within-subjects factors of memory assessment (error computed relative to studied and remembered locations) and testing round (memory testing rounds 1 and 2).

To examine whether cueing during the LDT impacted later memory, we primarily focused on assessing spatial error separately for cued and uncued object-location associations relative to each object’s placement during the immediate memory test (memory stability), which in the prior analysis was found to be a better predictor of delayed memory. A repeated measures ANOVA was performed on the average spatial error across trial types, using a within-subjects factor of cueing (cued/uncued associations) and testing round (Day 2 memory testing rounds 1 and 2). In addition to examining spatial error relative to object placement on Day 1, we also asked whether targeted memory reactivation influenced the stability of next-day representations in memory by computing the spatial distance between each object’s placement across the two rounds of memory testing on Day 2. We then compared this measure of stability across cued and uncued associations using an unpaired t-test. As a secondary goal, we also examined the influence of cueing on delayed object memory when measured relative to each object’s studied location, similar to prior work^19,44,54^. To do so, we performed a repeated measures ANOVA on averaged delayed memory relative to studied locations using a within-subjects factor of cueing (cued/uncued associations) and testing round (Day 2 memory testing rounds 1 and 2).

### Statistical Analysis: Mixture models

In addition to average measures of graded object-location memory, we also fit probabilistic mixture models to explicitly model object-location memory as the combination of a guessing process (resulting in uniformly distributed locations across the screen) and successful associative memory (resulting in locations centered around the true location with a Gaussian spread of error around this location). Similar to prior approaches modeling two-dimensional spatial locations^75^, two free parameters were estimated from the data: a memory success parameter which determined the mixing of the memory distribution versus the guessing distribution and the standard deviation of the two-dimensional Gaussian distribution used to model memory success, representing the precision of memory representations. The guessing distribution was determined by the guessing rate (1/the screen size), which is equal to 1/(1280*1024 pixels). The memory success distribution was modeled using mvnpdf in MATLAB, which estimates the probability density of a multi-variate normal distribution. The values of these free parameters were estimated using a maximum likelihood approach, such that the negative log likelihood of the mixture model was minimized using a constrained search via the fmincon MATLAB function. A constrained search was used in order to obtain plausible estimates of the free parameters (the memory success parameter was constrained to be between 0 and 1 and the standard deviations were constrained to be in the range of 1 – 800 pixels).

Using this mixture modeling approach, we asked whether delayed object placement was better predicted by its studied versus remembered location by fitting two models for each participant, using the true location for each association as either the studied or remember location (all trials from both rounds of delayed memory testing were used). We then compared memory precision (standard deviation of the fitted Gaussian distribution) for studied versus remembered locations using a paired t-test. Lastly, we examined how cueing impacted memory success by fitting mixture models in addition to graded memory (described above). As only eight trials were present for cued and uncued associations in each participant (not enough to obtain meaningful estimates of memory success and precision for each participant), we fit a mixture model at the group level using both cued and uncued memory measures. This process was performed separately considering memory relative to remembered locations and memory relative to studied locations. We then computed a binary measure of memory success for each trial and each type of memory assessment by estimating the probability of memory success based on the group level fitted Gaussian distribution and comparing this to the probability of belonging to the uniform guessing distribution. Each trial was then labeled as representing memory success if the Gaussian probability was greater than the guessing probability, and this memory success status was then averaged across trials for cued and uncued associations in each participant. Differential changes in memory success were assessed using a repeated-measures ANOVA with within-subject factors of cueing (cued and uncued associations) and testing round. Separate ANOVAs were performed to assess the influence of cueing on delayed memory success, considering memory relative to remembered locations as well as relative to studied locations.

### Statistical Analysis: Relation between LDT performance, vigilance, and cueing memory benefits

In addition to the overall impact of cueing on later memory, we asked whether participant’s behavioral state prior to cue presentation was related to the benefit of cueing on later memory. To do so, we measured the average reaction time (RT) during the arrow task (the ITI of the LDT) performed prior to each object re-exposure. As each object was presented three times during the LDT, we averaged the mean RT across these three ITI periods for each cued object. To obtain a normalized measure of RT that is comparable across participants, we subtracted the mean RT across ITIs in which cueing was not performed from the mean RT prior to each cued association. The same results were obtained when this RT measure was normalized by subtracting the mean RT across all ITIs in the LDT. The cueing benefit for each association was measured as the difference in error for that association from the mean error across all uncued associations for that participant, such that positive values reflect a greater reduction in error associated with cueing. We then used regression to assess the relationship between RT prior to cueing and the cueing-related benefit in later memory across all cued associations in all participants. Robust regression, as implemented in robustfit in MATLAB, was primarily used to assess this relationship. Robust regression seeks to minimize the influence of large outliers via an iterative process. This iterative process consists of fitting a standard ordinary least squares solution, and subsequently down-weighting data points with high residual error from the ordinary least squares solution in the next iteration of fitting an ordinary least squares solution. This process is repeated until convergence, defined as minimal changes in regression coefficients between iterations. The default parameters for robustfit were used in all analyses. To ensure that individual outlier trials were not influencing the relationship between RT during the ITI and the cueing memory benefit, we also excluded trials with a RT or cueing benefit that was two standard deviations from the mean and assess this relationship using standard ordinary least squares regression.

We also examined how cueing impacted subsequent performance on the LDT. To do so, we computed the average RT on lexical decision trials in which objects were and were not presented. A paired t-test was used to assess differences lexical decision RTs as a function of cueing. Similar to the prior analysis, we also asked whether trial-by-trial changes in RT after cue presentation were predictive of the impact of cueing on later memory. To do so, we used the same procedure as for the previous analysis, but used mean RTs on the lexical decision trial after cue presentation as opposed to RTs before cue presentation.

### Statistical Analysis: Relationships between initial learning and cueing memory benefits

We asked whether cueing benefits were related to initial learning/memory performance both at the level of individual participants and items^22,54^. At the subject level, we examined two measures related to initial learning and memory, the average error of object placement during immediate memory testing and the mean number of training blocks needed to reach learning criterion (averaged across associations within a participant). We first examined the correlation between these measures. Since they were highly correlated, we performed principal components analysis (PCA) to extract a factor that represented their shared variance. The 1^st^ principal component was highly correlated with both measures (r = .94 with immediate memory, r = .91 with number of blocks to reach training criterion). We then used this 1^st^ principal component as a factor representing initial learning/memory and assessed the relationship between this measure and the benefit in memory stability associated with cueing (cued minus uncued distance from immediate to delayed testing, averaged across testing rounds during the delayed memory test) across participants using robust regression. To examine whether cueing benefits were related to initial memory at the level of individual associations, we performed a split half analysis on individual trials for each participant based on initial memory, separately for cued and uncued associations. We then performed a repeated measures ANOVA with factors of initial memory (low/high initial error), cueing (cued/uncued associations) and rounds of delayed testing. Follow up t-tests were performed to examine cueing effects (differences in cued versus uncued memory stability) separately for low and high error associations.

### Statistical Analysis: Cue discrimination task

To assess whether participants could distinguish between cued versus uncued objects, we examined the distribution of responses on the cue discrimination task for cued and uncued objects using a repeated measures ANOVA with factors of cueing and response option (high confident cued, low confident cued, low confident uncued, high confident uncued). We then performed follow up t-tests to assess differences between the proportion of cued versus uncued trials for each response option. Our primary question was whether cueing benefits were driven by explicit knowledge of which items were cued versus not cued. We first addressed this question at the level of individual participants, using a chi-squared test to assess whether each participant showed above-chance discrimination of cued versus uncued objects. We computed a chi-squared statistic based on the number of trials labeled as high confident cued versus the other response options and evaluated significance of the chi-squared statistic relative to a chi distribution. Participants were labeled as showing above-chance cue discrimination of their p-value for significance testing showed a trend (was below .1), while all other participants were labeled as non-cue-discriminators. An unpaired t-test was used examine whether cueing benefits (cued minus uncued error averaged across delayed testing rounds) differed for cue discriminators versus non cue discriminators. Lastly, we examined whether cueing benefits were still present for items labeled as uncued. To do so, we computed the average memory stability for associations, excluding those in which the object was labeled as cued in the cue discrimination task. We also performed the complementary analysis and examined memory stability for associations in which the corresponding object was labeled as cued. For each of these analyses, some participants did not contribute any trials when particular trial types were excluded, resulting in smaller sample sizes (N = 21 for associations labeled uncued, N = 17 for associations labeled cued).

## ACKNOWLEDGEMENTS

We thank Anuya Patil for assistance with data collection and constructive comments and feedback provided by anonymous reviewers. This work was supported by Dart Neuroscience and the National Institutes of Health (MH074692) to L.D.

## AUTHOR CONTRIBUTIONS

All authors designed the experiment and wrote the paper. A.T. analyzed the data.

